# Effects of Hydrolyzed Collagen as a Dietary Supplement on Fibroblast Activation: A Systematic Review

**DOI:** 10.1101/2024.03.06.583701

**Authors:** Pedro Augusto Querido Inacio, Felipe Augusto Chalupe, Rodolfo P Vieira

**Affiliations:** Evangelical University of Goias (Unievangelica), Laboratory of Pulmonary and Exercise Immunology (LABPEI), Avenida Universitária Km 3,5, Anápolis – GO, 75083- 515, Brazil; Peptech Colagen from Brazil, 1500 North Halsted Street - Floor 2, Chicago – IL, 60642-2517, USA

**Keywords:** Low molecular weight collagen peptides, Fibroblast, Hydrolyzed Collagen, Collagen supplementation

## Abstract

**Purpose:** Our objective was to conduct a systematic review of the effects of hydrolyzed collagen supplementation on the proliferation and activation of fibroblasts.

**Methods:** The search was conducted for journals that published articles in the English language, peer-reviewed, meeting the following criteria: a) randomized clinical trials, b) randomized studies in animals or humans, c) in vitro studies, d) studies using hydrolyzed collagens or collagen peptides, d) studies assessing alterations on fibroblasts as primary or secondary outcome. We utilized the main journal databases PUBMED/WEB OF SCIENCE and ongoing reviews by PROSPERO. For bias risk and methodological quality, we used an adaptation of the Downs and Black checklist. Our review followed the PRISMA checklist, conducted from February 2024 to the first week of March 2024, by two independent researchers (P.A.Q.I. and R.P.V.).

**Results:** Eleven studies were included in this review, where our findings reinforce the notion that hydrolyzed collagens or collagen peptides at concentrations of 50-500 μg/ml are sufficient to stimulate fibroblasts in human and animal tissues without inducing toxicity. Different enzymatic processes may confer distinct biological properties to collagens, allowing for scenarios favoring fibroblast promotion or antioxidant effects. Lastly, collagens with lower molecular weights exhibit greater bioavailability to adjacent tissues.

**Conclusion:** Hydrolyzed collagens or collagen peptides with molecular sizes ranging from <3 - 3000 KDa promote the stimulation of fibroblasts in human tissues.

## Introduction

Collagen is the primary protein composing the extracellular matrix, formed by a chain of specific amino acids. In its primary form, collagen exhibits a triple-helical structure (H. Wang, 2021). The triple helical structure of collagen molecules is constituted by three α chains, characterized by repeating peptide triplets with the sequence glycine-X-Y. While X and Y represent any amino acid, they typically correspond to proline and hydroxyproline, respectively. Adjacent to the triple helical regions are non-glycine-X-Y segments known as non-collagenous domains or Col domains. These regions often incorporate identifiable peptide modules found in other extracellular matrix molecules (Gordon et al., 2010).

In addition, collagen family encompass 28 types of collagens which are distributed in different proportions in all organs and systems in the human body, beyond to constitute an essential component of cellular membranes (S. Ricard-Blum, 2011). Although almost of collagens are presented in filamentous or fiber forms, few forms are also presented in soluble structure (S. Ricard-Blum, 2011). In this way, collagens fulfill structural functions and play crucial roles in determining the mechanical characteristics, arrangement, and form of tissues. They engage with cells through various receptor groups, influencing their growth, movement, and specialization. Certain types of collagens exhibit limited distribution within tissues, indicating specialized functions tailored to specific biological processes (S. Ricard-Blum, 2011). The proper functioning of tissues relies on the accurate assembly of molecular aggregates integrated into the matrix. This review underscores key structural attributes of collagen types I-XXVIII (Gordon et al., 2010).

Collagens stand as the predominant proteins in mammals, constituting approximately 30% of the total protein mass. The unveiling of collagen II by Miller and Matukas in 1969 marked a pivotal moment, leading to the subsequent identification of 26 additional collagen types. Advances in molecular biology and gene cloning techniques have greatly expedited the pace of discovery in this field (Gordon et al., 2010).

Beyond their physiological and pathophysiological roles, the development of collagen-based proteins and peptides, are expanding rapidly (Xu et al., 2023). Through a process of heating, the collagen molecule undergoes enzymatic denaturation, partly through the hydrolysis of peptide bonds, resulting in its gelatinous form, commercially known as hydrolyzed collagen or collagen peptides (Figueres Juher & Bases Perez, 2015; Zague, 2008; Zhuang, Hou, Zhao, Zhang, & Li, 2009). Its production occurs through animal skin, bones, connective tissues (tendons), or the skin and scales of saltwater and freshwater fish (Pati et al., 2012). The growing interest of pharmaceutical and cosmetic industries is due to the evidenced effects, mainly on the regeneration of wrinkled cutaneous tissue and dermal hydration, seemingly resulting from the recovery of degraded collagen fibers (Pyun et al., 2012).

Hydrolyzed collagen, also known as collagen peptides, is a popular dietary supplement widely used for its potential health benefits, including improved skin elasticity, joint health, and bone density. Industrial production of hydrolyzed collagen involves several key techniques to extract and process collagen from animal sources, primarily bovine (cow) or porcine (pig) hides, bones, or fish scales. So, the industrial techniques used in the production of commercial hydrolyzed collagen are briefly described as follow:

Extraction of Collagen: The first step involves the extraction of collagen from animal tissues. This is typically done through acid or alkaline treatment to break down the connective tissue and release the collagen fibers. Various studies detail the optimization of extraction parameters such as temperature, pH, and extraction time to maximize collagen yield (Bi et al., 2019).

Purification: Once extracted, the collagen undergoes purification processes to remove impurities such as fats, proteins, and minerals. Techniques like filtration, centrifugation, and precipitation are employed for this purpose (Bi et al., 2019).

Hydrolysis: The purified collagen is then subjected to enzymatic hydrolysis, where enzymes such as proteases break down the collagen molecules into smaller peptides. This process increases the bioavailability and solubility of collagen peptides, making them easier to digest and absorb in the body (Vidal et al., 2020).

Fractionation: Fractionation techniques are sometimes employed to isolate collagen peptides of specific molecular weights or sizes, as different sizes may exhibit different bioactive properties. Fractionation methods include chromatography and ultrafiltration (Nekliudov et al., 2003).

Drying and Packaging: The hydrolyzed collagen solution is then concentrated and dried to obtain a powder form suitable for use as a dietary supplement. Spray drying is a commonly used technique for this purpose, as it allows for rapid drying while preserving the nutritional and functional properties of collagen peptides. The dried powder is then packaged in airtight containers to maintain its quality and shelf life (Vargas-Muñoz and Kurozawa et al., 2020).

Quality Control: Throughout the production process, quality control measures are implemented to ensure the safety, purity, and consistency of the final product. This includes testing for contaminants, such as heavy metals and pathogens, as well as monitoring key parameters like collagen content, amino acid profile, and molecular weight distribution (López-Morales et al., 2019).

Regulatory Compliance: Industrial production of hydrolyzed collagen must comply with regulatory standards set by organizations such as the Food and Drug Administration (FDA) in the United States or the European Food Safety Authority (EFSA) in Europe. This includes adherence to good manufacturing practices (GMP) and labeling requirements for dietary supplements (Burdok and Carabin., 2004).

In terms of health implications, collagen supplementation has been implicated in elderly sarcopenic populations, wound healing, and degenerative processes related to tendon and bone structures (H. Wang, 2021). New findings also suggest that hydrolyzed collagen possesses properties to reduce inflammatory markers and fibroblast proliferation. This emerging research showing this relationship is justified by the mechanism of action in which collagens are formed by fibroblasts (Brandao-Rangel et al., 2022), supporting the notion that collagen appears to stimulate fibroblast proliferation. Animal studies and cell culture experiments corroborate with this rationale (Chotphruethipong, Sukketsiri, Aluko, Sae-Leaw, & Benjakul, 2021; Offengenden, Chakrabarti, & Wu, 2018).

Fibroblasts were first introduced to the scientific community in 1858 by the German pathologist Rudolf Virchow, and subsequently, the properties of this discovery turned towards their role in tissue repair, implicating fibroblasts in the production of new connective tissues (Molenaar, 2003). Broadly, fibroblasts are known as mesenchymal cells, exhibiting various functions such as tissue repair agents in various pathological scenarios, target cells, and modulators for immune functions, secretion of metabolites and signaling factors to neighboring cells, as well as regulation of metabolism in various organs, and extracellular matrix remodeling (Plikus et al., 2021).

Based on these findings, we investigated how supplementation with hydrolyzed collagen peptides can induce the proliferation and activation of fibroblasts, which are the cells capable of synthesizing collagen in various organs and tissues. The justification for this literature review stems from the lack of a review that explains the studies corroborating the idea of “hydrolyzed collagen inducing fibroblast stimulation.” The objective of this review is to present possible evidence that brings to light fibroblast proliferation through the ingestion or application of hydrolyzed collagen peptides and their reduced forms, to highlight limitations, and to create a more solid database for the mechanisms adjacent to the use of hydrolyzed collagen.

## Methods

### Inclusion and Exclusion Criteria

Our analyses were limited to journals that published articles in the English language, peer-reviewed, meeting the following criteria: a) randomized clinical trials, Nb) randomized studies in animals or humans, c) in vitro studies, d) studies using hydrolyzed collagens or collagen peptides, d) studies assessing alterations on fibroblasts as primary or secondary outcome. Review articles, brief reviews, opinion articles, case reports, or observational studies, as well as research utilizing forms of non-hydrolyzed collagens, were considered ineligible for the construction of this review.

### Search Strategy

The search for articles, as well as the construction of this review, was conducted in accordance with the Preferred Reporting Items for Systematic Reviews and Meta-Analyses (PRISMA) guidelines (Moher, Liberati, Tetzlaff, Altman, & Group, 2009). Our primary search was performed in the main databases PubMed/MEDLINE, Web of Science/Clarivate, which publish articles in the English language. We also consulted the ongoing review database (PROSPERO). The search was conducted from February 2024 to the first week of March 2024. The following syntax was used for the search: hydrolyzed collagen peptides AND fibroblast, low molecular weight collagen peptides AND fibroblast. Articles that included any of the following terms in their title or abstract: Collagen hydrolyzed, fibroblast, Low-molecular-weight collagen, were included for further analysis. From these selected articles, the reference lists were subsequently consulted to identify possible additional publications (Greenhalgh & Peacock, 2005). To avoid selection bias, the search was conducted by two researchers who independently analyzed the articles (P.I and R.P.V). If any article was deemed divergent for inclusion and exclusion, a third researcher (F.A.C) was consulted, with the final decision resting with them.

### Methodological Quality

We assessed the methodological quality of the included studies using the Downs and Black checklist with 27 items (Downs & Black, 1998). This checklist addresses different aspects of study design, including reporting (Items 1–10), external validity (Items 11–13), internal validity (Items 14–26), and statistical power (Item 27). Given the specificity of the included studies, we used the checklist as a reference and listed 10 items to be considered to evaluate the methodological quality of the studies. If the item was present, a value of 1 was assigned, otherwise a value of 0, similar to the Downs and Black checklist. Studies were classified as good quality if they scored 8-10, moderate quality if they scored 6-7, and poor quality if they scored 5 or below. The studies were independently evaluated by 2 reviewers (P.I and R.P.V).

## Results

From the studies searched in the databases, we obtained a total of 220 articles, of which 184 articles were listed for the initial analysis. However, these were subsequently excluded due to ineligibilities identified by automation tools, resulting in the exclusion of a total of 172 documents, including duplicates (n=8) and articles published in languages other than English (n=4), leaving 36 articles. After summarization, those that did not meet criteria a); b); c); d), or were not considered relevant for the review construct, in agreement between the two researchers (P.I and R.P.V), were excluded (n=21), totaling 15. Three articles were evaluated by a third researcher (F.A.C), who made the final decision regarding their inclusion or exclusion in the present review. Two articles proceeded for analysis, and one document was excluded. Subsequently, the articles were read in full, and the references were re-evaluated to retrieve possible documents not found in the initial data searches. During the full-text reading of the articles, three articles were excluded: one as it was a review, and the other two documents did not demonstrate in their results the effects of hydrolyzed collagen on fibroblast proliferation. Eleven articles were included in the final review. Figure 1 presents the flowchart of the literature search. In summary, the studies included for analysis are as follows.

**Figure 1.**
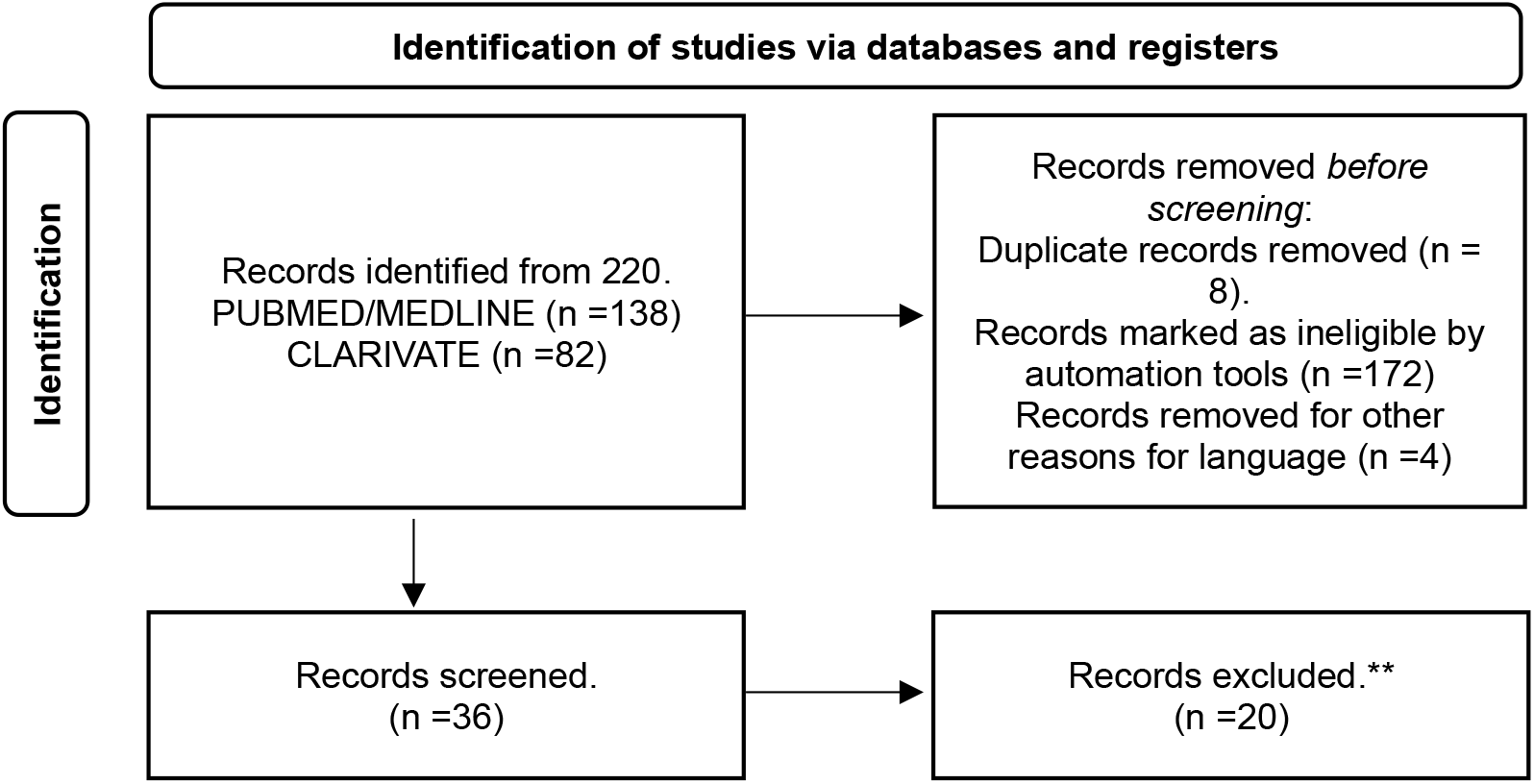
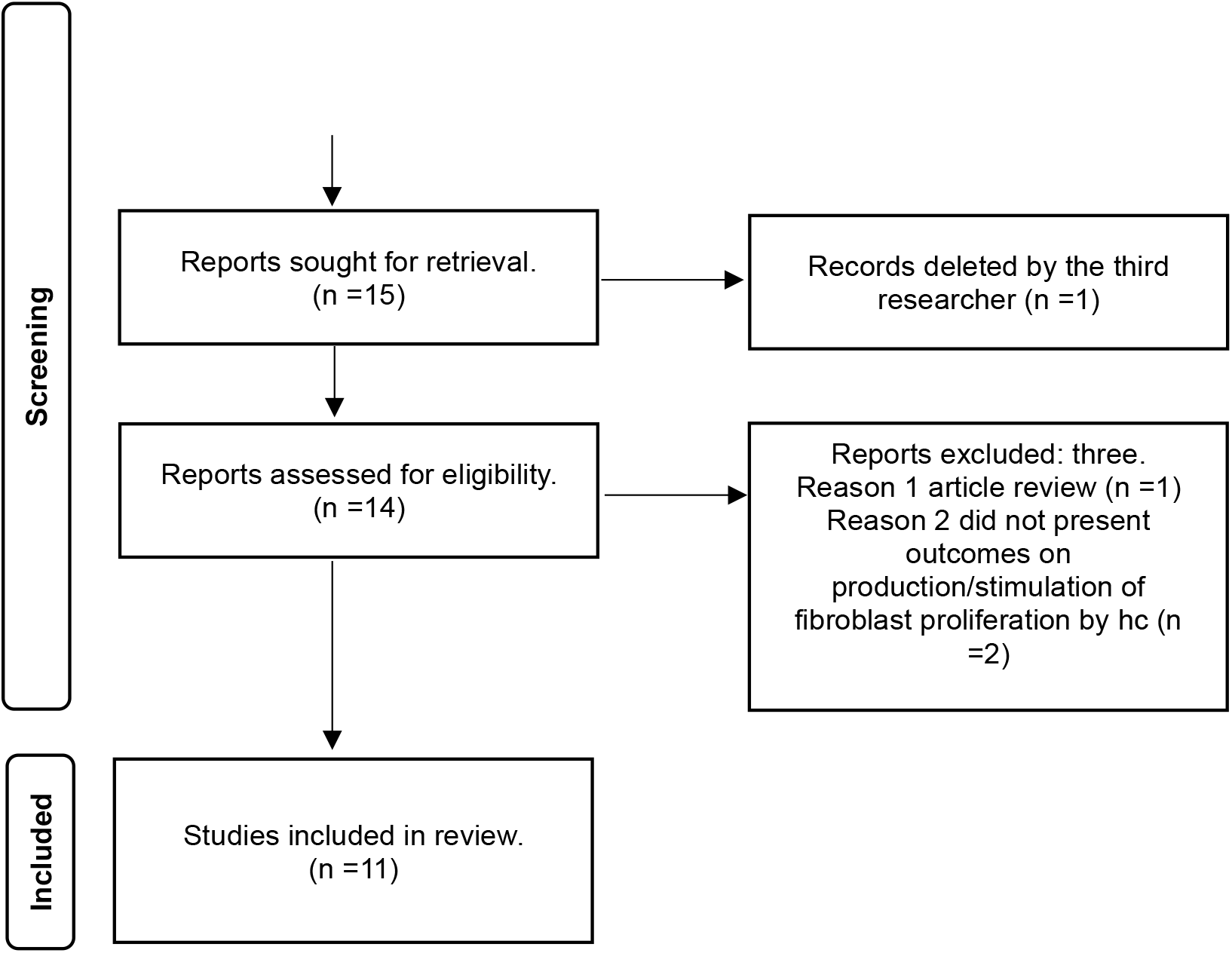
Flowchart of the search process.

### Study Coding

The studies were individually read and coded by two researchers (P.I and R.P.V) using the PICO tool for the following variables: (a) Population, (b) Intervention, (c) Comparison/Control, (d) Outcomes. Population refers to the type of hydrolyzed collagen used and, if reported, its molecular weight in kilodalton KDa. Intervention on fibroblasts identifies the types of fibroblastic cells used, whether human or animal. For comparison and control, we listed the controls used as well as the dosages. Lastly, Outcomes present the results exclusively regarding fibroblast cell proliferation. Table 1 presents an overview of the studies that met all inclusion criteria.

**Table 1.**
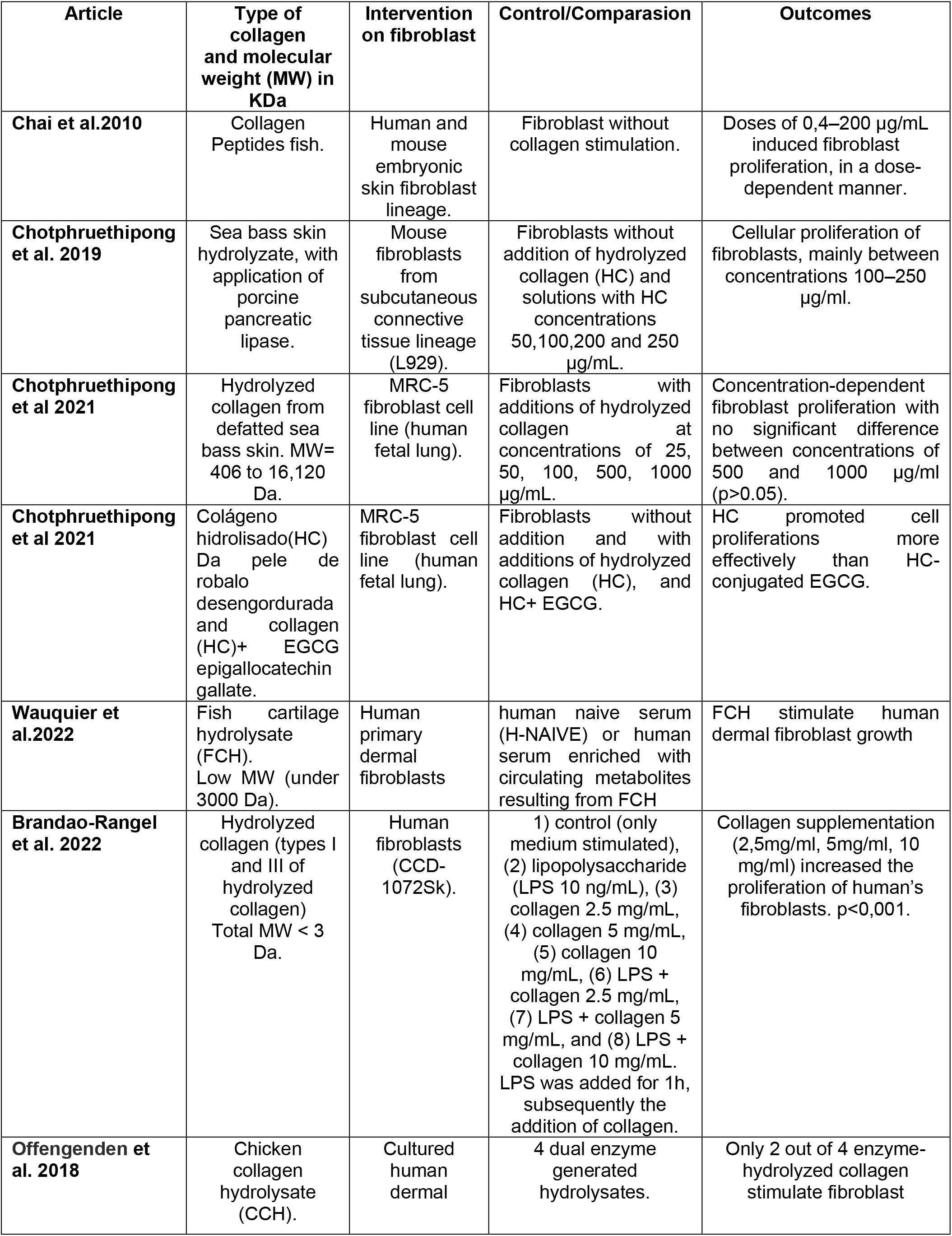
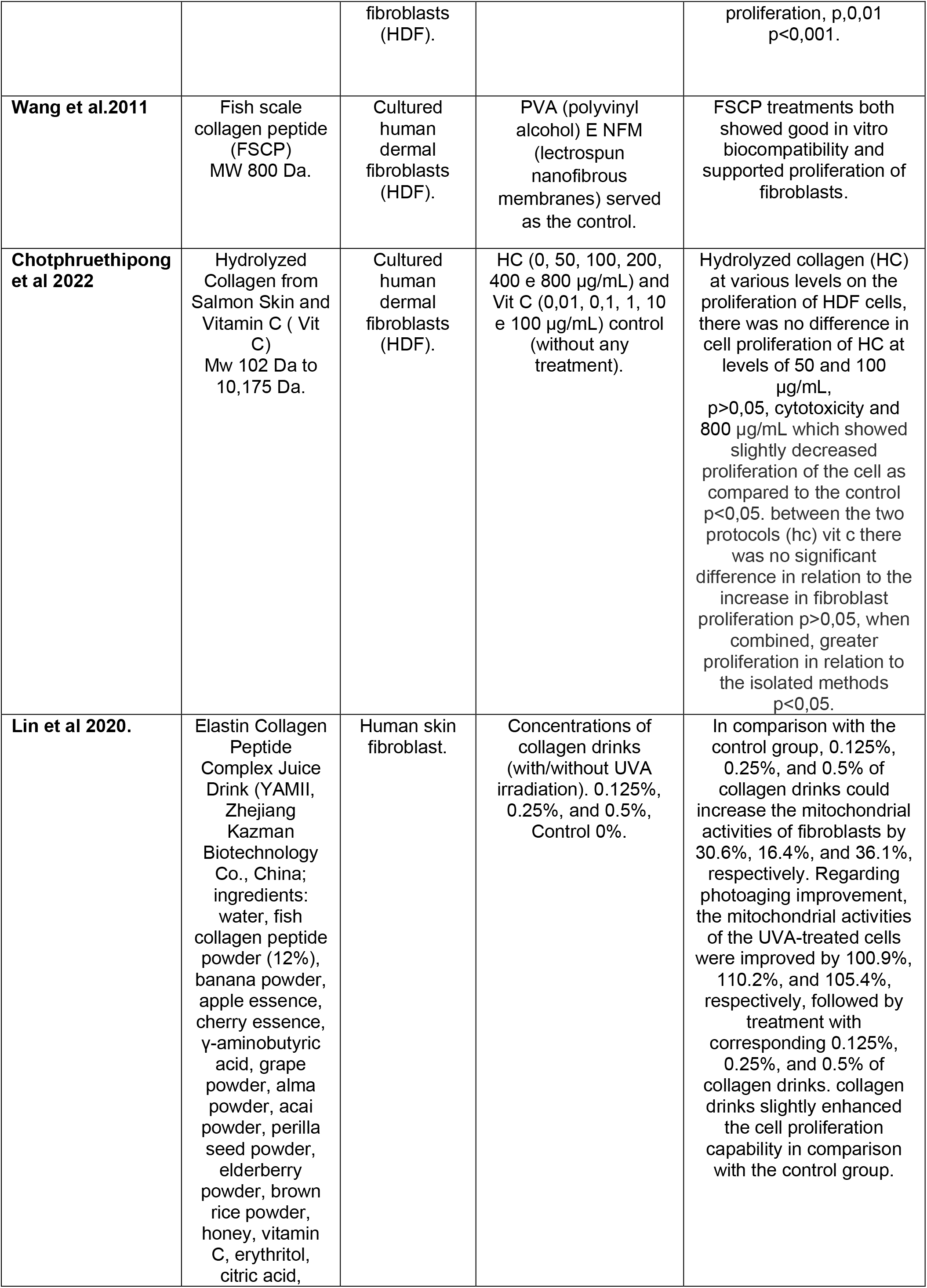
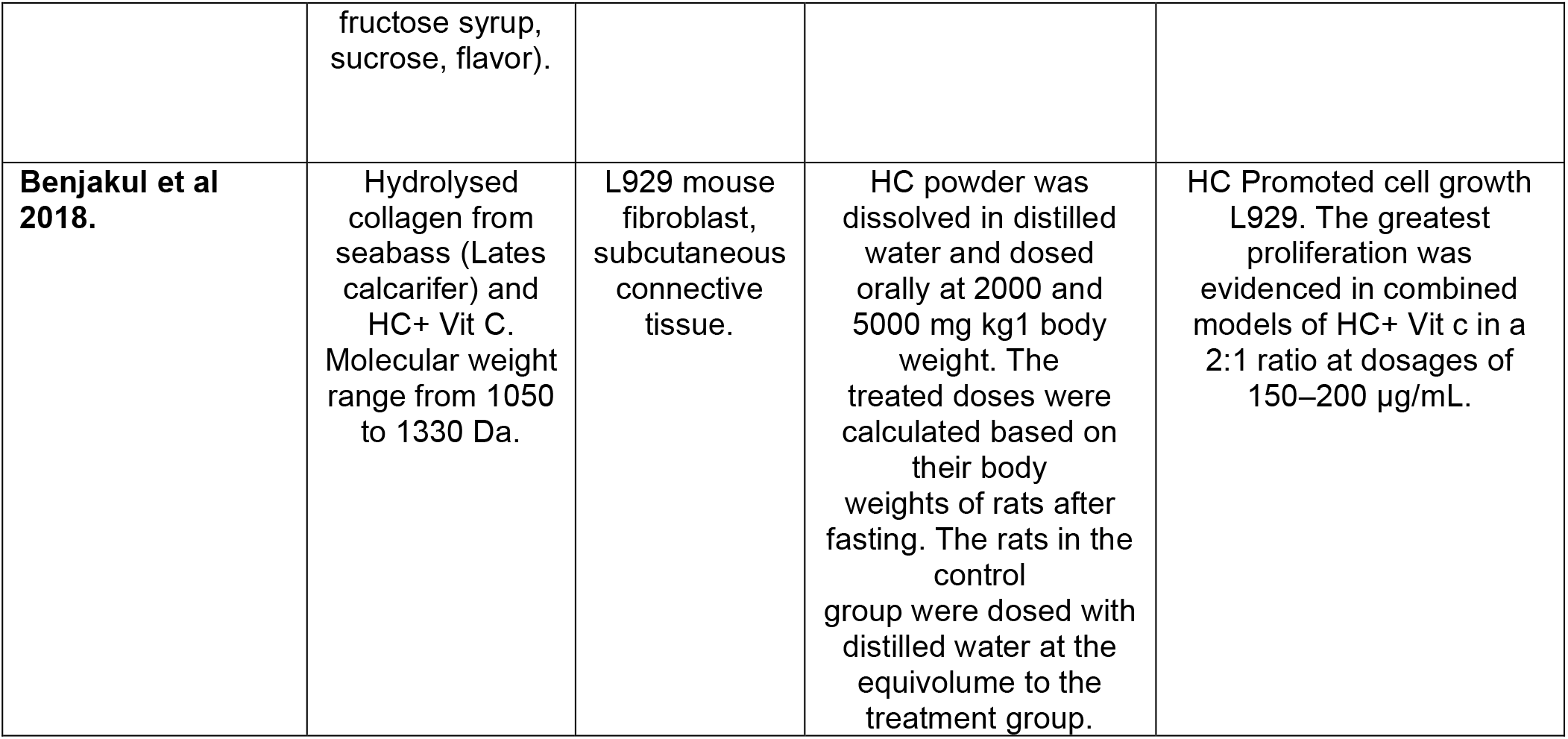
Articles included for review and their outcomes.

**Table 1**

### Article Retrieval

Throughout the process of reading and summarizing the articles, their references and citations were independently reviewed to find articles that were not reached in our initial searches. However, no articles that had not already been identified during the review process were found. From all references, the researcher (P.I) listed one document that addressed the outcome of fibroblast proliferation by collagens. Upon reading the document in its entirety, it became evident that the document aimed to evaluate the effects of collagen that did not undergo hydrolysis processes, thus disqualifying it as hydrolyzed collagen or collagen peptides. Therefore, it was not included.

### Characteristics of the Studies

Eleven studies were included in this review (Benjakul, Karnjanapratum, & Visessanguan, 2018; Brandao-Rangel et al., 2022; Chai et al., 2010; Chotphruethipong, Aluko, & Benjakul, 2019; Chotphruethipong et al., 2022; Chotphruethipong, Sukketsiri, Aluko, et al., 2021; Chotphruethipong, Sukketsiri, Battino, & Benjakul, 2021; Lin et al., 2020; Offengenden et al., 2018; Y. Wang, Zhang, Zhang, & Li, 2011; Wauquier et al., 2022) for the outcome of stimulation or proliferation of fibroblasts by the application or ingestion of hydrolyzed collagen supplements or collagen peptides. Three studies used rat fibroblast cell lines as the intervention method, while the remaining articles used human fibroblast cells. The types of collagens used varied among the selected articles, including animal, fish, and chicken origins, with one study utilizing a collagen-enriched beverage (Lin et al., 2020), and two studies combining hydrolyzed collagen with vitamin C applications (Benjakul et al., 2018; Chotphruethipong et al., 2022). When reported, the molecular weight of all hydrolyzed collagens was stated in Daltons, with all studies presenting low molecular weights, with the lowest being reported in the work of Rangel et al., 2022 (See Table 1). The studies used control cells without interventions or with additions of various concentrations of hydrolyzed collagen (25, 50, 100, 500, 1000 μg/mL).

### Methodological Quality

The quality of each study was assessed by two researchers (P.I and R.P.V), and in case of any discrepancies regarding quality parameters, a third researcher was consulted to ensure agreement. The scale used was adapted from the Downs and Black checklist, listing 10 points as quality criteria, of which eight studies were considered good quality, and three studies were of moderate quality (See Table 2).

**Table 2.**
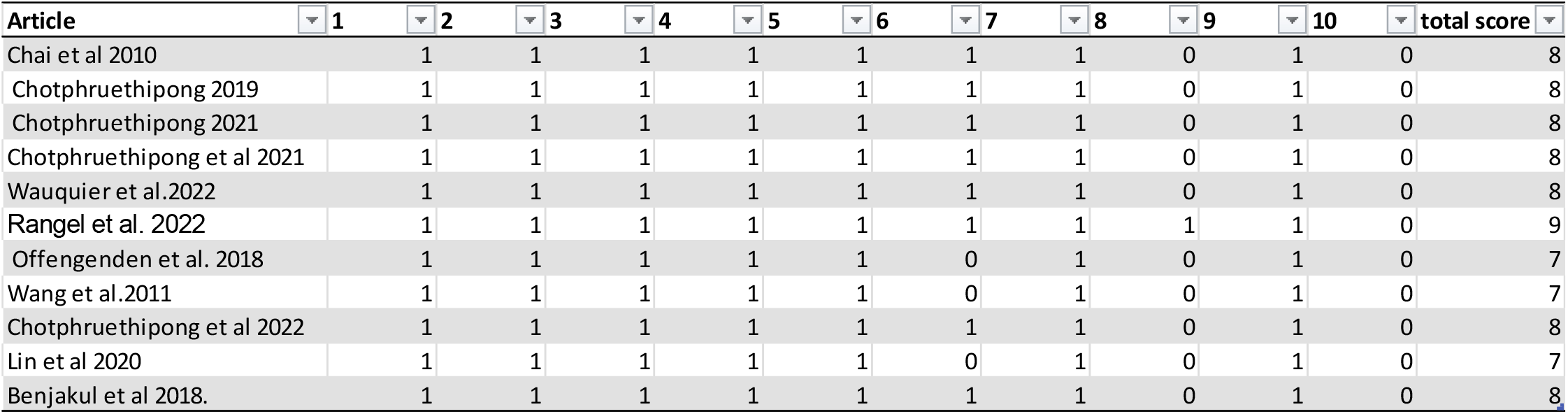
Methodological quality criteria.

**Table 2**

## Discussion

The results of this systematic review reinforce the concept of “hydrolyzed collagen inducing fibroblast stimulation.” This effect appears to be mediated by the ingestion of collagen peptides or hydrolyzed collagens. The studies presented here provide a robust body of evidence demonstrating fibroblast cell proliferation through the use of hydrolyzed collagens without alterations in cellular toxicity. This implies that hydrolyzed collagen may have outcomes beyond those already evidenced.

Hydrolyzed collagen supplementation is widely used for aesthetic purposes, primarily to delay skin aging. Part of its effect is attributed to stimulating extracellular matrix proteins, which are directly regulated by dermal fibroblasts (F.D). As shown in our results, hydrolyzed collagens stimulate the specific growth of F.D, especially those with lower molecular weights and high proline/hydroxyproline content.

A determinant factor reported in our review leads us to rationalize that molecular weight is directly related to better solubility. Collagen peptides rich in pro-Hdx exhibit significant bioavailability. Peptides with lower molecular weights have properties to reach deeper dermal layers, thus stimulating better tissue repair. Despite not being the focus of this review, it is important to note that, in addition to increasing fibroblasts, hydrolyzed collagens are gaining traction as potential antioxidant supplements. Some studies included here sought to combine hydrolyzed collagens with other types of supplements known for their positive effects on oxidative scenarios, such as vitamin C.

Among the findings, it was also noted that depending on the enzymatic processes on collagens, different hydrolyzed collagens obtain distinct characteristics. Some seem to prefer antioxidant and inflammatory situations, while others induce fibroblast proliferation in human skin. However, this result is presented by only one study, warranting further investigation to identify if indeed specific types of hydrolyzed collagens are tailored to specific fibroblast applications.

According to our findings, hydrolyzed collagens from different sources and with different molecular weights, yet all considered low-weight, ranging between <3-3000 Da, are capable of inducing fibroblasts in human and animal skin tissues.

In addition to the findings presented here, this review also serves an important role in scientific production, encouraging new studies to evaluate the effects of hydrolyzed collagens on fibroblasts in relation to the tested dosages. Most studies indexed here used dosages with different concentrations of collagens in fibroblasts to identify minimum and maximum absorption values and possible toxicity. Summarizing all presented results, concentrations of 50-500 μg/ml are sufficient for fibroblast proliferation without toxicity implications. Concentrations of 800 μg/ml showed toxicity levels in Cultured Human Dermal Fibroblasts (HDF). However, this result should be analyzed cautiously since in MRC-5 fibroblast cell line (human fetal lung) tissues, doses of 1000 μg/ml did not create such a scenario, and among lower doses compared to higher doses, no better dose responses were evident. With these reviewed findings, new research directions do not require further investigation into solutions with different concentrations when the goal is fibroblast stimulation.

### Limitations and Future Directions

This review focused on only one outcome regarding hydrolyzed collagens, while others such as toxicity and inflammatory levels are recommended for readers to explore in the selected articles. Our aim is to facilitate new research regarding the use of collagens and fibroblasts, as until the present moment, no review with such objectives has been evidenced. Our work is not without limitations, for example, the methodological quality of the listed articles underwent an adaptation process. We encourage future, more robust checklists that include in vitro studies to be developed to facilitate methodological processes for cellular study reviews. We chose to limit our searches to articles published in English and peer-reviewed, which may omit data from other sources that would be relevant for a better understanding of the presented findings. However, this choice aims to narrow down the findings to studies with minimal bias and good quality. Finally, this review creates a possible research scenario regarding the specific biological properties of each hydrolyzed collagen, and in the future, identify if specific enzymatic processes substantially alter the action of hydrolyzed collagen on fibroblasts.

## Conclusion

In conclusion, our review highlights the crucial role of hydrolyzed collagens and collagen peptides in stimulating and proliferating fibroblast cells. New studies aiming to explore this scenario may consider using dosages ranging between concentrations of 50-500 μg/ml of hydrolyzed collagen. Furthermore, investigations into enzymatic changes during the collagen hydrolysis process should be conducted to identify if these alterations affect the biological properties, thus changing their effects on fibroblast stimulation.

## ATTACHMENTS

### Methodological quality criteria adapted from the checklist

1. Was the study hypothesis/goal/objective clearly described?
2. Were the main results clearly measured and described in the Introduction or Methods section?
3. Were the characteristics of the cell types used and controls described?
4. Were the interventions of interest clearly described? Treatments and placebos (when relevant) to be compared should be clearly described.
5. Was a list of key confounding factors provided, along with how cellular environments and processes were managed and described clearly?
6. Were the main conclusions of the study clearly described? Simple outcome data (including denominators and numerators) should be reported for all main findings so that the reader can verify the main analyses and conclusions. (This question does not cover statistical test data, which are considered below).
7. For non-normally distributed data, was the interquartile range of results reported? For normally distributed data, standard error, standard deviation, or confidence intervals should be reported.
8. Did the authors clearly report limitations of their findings? If yes, consider 1; if not, consider 0.
9. Were appropriate statistical tests used to assess the main results? The statistical techniques used should be appropriate to the data.
10. Were actual probability values reported (e.g., 0.035 instead of <0.05) for the main results, except where the probability value is less than 0.001?

